# Endogenous p53 expression in human and mouse is not regulated by its 3′UTR

**DOI:** 10.1101/2020.11.23.394197

**Authors:** Sibylle Mitschka, Christine Mayr

## Abstract

The *TP53* gene encodes the tumor suppressor p53, which is functionally inactivated in many human cancers. Numerous studies found that overexpression of specific microRNAs or RNA-binding proteins can alter p53 expression through binding to *cis*-regulatory elements in the *TP53* 3′ untranslated region (3′UTR). Although these studies suggested that 3′UTR-mediated p53 expression regulation could play a role in tumorigenesis or could be exploited for therapeutic purposes, they did not investigate post-transcriptional regulation of the native *TP53* gene. We used CRISPR/Cas9 to delete the human and mouse p53 3′UTRs while preserving endogenous mRNA processing. This revealed that the endogenous 3′UTR is not involved in regulating p53 mRNA or protein expression neither in steady state nor after genotoxic stress. As we were able to confirm the previously observed repressive effects of the isolated 3′UTR in reporter assays, our data highlight the importance of genetic models in the validation of post-transcriptional gene regulatory effects.

## Introduction

The transcription factor p53 coordinates the cellular stress response. p53 regulates expression of genes involved in cell cycle control, DNA repair, apoptosis, metabolism, and cell differentiation (Kastenhuber and Lowe, 2017). Reduced levels or insufficient p53 activity are major risk factors for the development of cancer and more than half of all human cancers exhibit diminished p53 expression or function (Kastenhuber and Lowe, 2017). In contrast, hyperactive p53 has been linked to impaired wound healing, obesity and accelerated aging (Rufini et al., 2013). These phenomena highlight the importance of p53 protein abundance and activity regulation in human health. p53 protein abundance is primarily controlled by a regulatory feedback loop involving the ubiquitin ligase MDM2. In addition, post-translational modifications of p53 and cofactor recruitment regulate its transcriptional activity (Hafner et al., 2019).

The 3′UTR of the *TP53* mRNA is another widely studied element of p53 expression regulation. Apart from facilitating pre-mRNA processing, 3′UTRs can also recruit microRNAs (miRNAs), RNA-binding proteins, and lncRNAs to modulate mRNA stability and protein translation (Tian and Manley, 2017; Mayr, 2019). The human p53 3′UTR contains experimentally characterized binding sites for 23 miRNAs, one lncRNA, and six RNA-binding proteins (Haronikova et al., 2019). A large number of experiments demonstrated the repressive nature of the *TP53* 3′UTR using reporter assays under steady state conditions (Table 1) (Haronikova et al., 2019). In addition, the *TP53* 3′UTR was shown to facilitate an increase in p53 translation after genotoxic stress (Fu and Benchimol, 1997; Mazan-Mamczarz et al., 2003; Chen and Kastan, 2010). This large body of work strongly suggested that miRNAs and RNA-binding proteins prevent p53 hyperactivation under normal conditions and induce p53 protein translation after exposure to genotoxic stress (Fu and Benchimol, 1997; Mazan-Mamczarz et al., 2003; Chen and Kastan, 2010). However, these claims have not been investigated under native conditions using the endogenous *TP53* mRNA.

**Table 1.** Previously reported evidence of miRNAs, lncRNAs, and RNA-binding proteins that target the p53 3′UTR.

| Interactors of the human *TP53* mRNA mapping to the last exon |  |  |  |  |  |
| --- | --- | --- | --- | --- | --- |
| Name | Type | Binding region (NM_000546.6) | Affected in dUTR allele? | Experiments | References (PMID) |
| miR-1228-3p | miRNA | 1422-1428 | yes | LRA, RT-qPCR, IHC, WB | 25422913 |
| miR-125a-5p | miRNA | 2044-2063 | yes | LRA, NB, RT-qPCR, WB | 19818772 |
| miR-125b-5p | miRNA | 2043-2064 | yes | LRA, ISH, RT-qPCR, WB | 19293287, 21935352, 27592685 |
| miR-1285-3p | miRNA | 2113-2134 | yes | LRA, RT-qPCR, WB | 20417621 |
| miR-150-5p | miRNA | 1568-1580 | yes | LRA, WB | 23747308 |
| miR-151a-5p | miRNA | 2304-2325 | yes | LRA, ChIP-seq, RT-qPCR, WB | 27191259 |
| miR-200a-3p | miRNA | 2269-2291 | yes | LRA, WB | 23144891 |
| miR-24-3p | miRNA | 2352-2374 | yes | LRA, IHC, RT-qPCR, WB | 27780140 |
| miR-25-3p | miRNA | 1401-1423 | yes | LRA, RT-qPCR, WB | 20935678 |
| miR-30d-5p | miRNA | 1596-1618 | yes | LRA, RT-qPCR, WB | 20935678 |
| miR-375 | miRNA | 1462-1483 | yes | LRA, Flow, RT-qPCR, WB, IF | 23835407 |
| miR-663a | miRNA | 1260-1281 | no (in CDS) | LRA | 27105517 |
| miR-504 | miRNA | 2059-2066, 2387-2395 | yes, no | LRA, RT-qPCR, WB | 20542001 |
| miR-92 | miRNA | 1417-1422 | yes | LRA, WB | 21112562 |
| miR-141 | miRNA | 2285-2290 | yes | LRA, WB | 21112562 |
| miR-638 | miRNA | 1381-1404 | yes | LRA, WB, IP | 25088422 |
| miR-3151 | miRNA | 1337-1354 | yes | LRA, WB, RT-qPCR | 24736457 |
| miR-33 | miRNA | 1957-1980 | yes | LRA, WB | 20703086 |
| miR-380-5p | miRNA | 1909-1936, 1943-1974 | yes, yes | LRA, WB | 20871609 |
| miR-19b | miRNA | 1712-1734 | yes | LRA, WB | 24742936 |
| miR-15a | miRNA | 2394-2414 | no | LRA, WB | 21205967 |
| miR-16 | miRNA | 2394-2415 | no | LRA, WB | 21205967 |
| miR-584 | miRNA | 1263-1284 | no (in CDS) | LRA, WB, IP | 25088422 |
| WIG1 | RBP | 2064-2106 | yes | LRA, IP, RT-qPCR | 19805223 |
| PARN | RBP | 2071-2102 | yes | LRA, EMSA, IP, RT-qPCR | 23401530 |
| CPEB1 | RBP | 2458-2500 | no | IP, RT-PCR | 19141477 |
| RBM38 (RNPC1) | RBP | 2064-2106 | yes | EMSA, IP, RT-PCR, Polysome gradient | 21764855, 24142875, 25823026 |
| RBM24 | RBP | 2064-2106 | yes | LRA, EMSA, IP, RT-qPCR | 29358667 |
| HUR | RBP | 2064-2106, 2393-2412, 2458-2505 | yes, yes, no | LRA, EMSA, WB, RT-qPCR | 12821781, 14517280, 16690610, 18680106 |
| 7SL | lncRNA | 2107-2149, 2194-2240, 2269-2301, 2307-2362 | yes, yes, yes, yes | LRA, IP, WB | 25123665 |

| Interactors of the murine *Trp53* mRNA mapping to the last exon |  |  |  |  |  |
| --- | --- | --- | --- | --- | --- |
| Name | Type | Binding region (NM_011640.3) | Affected in dUTR allele? | Experiments | References (PMID) |
| miR-92a-3p | miRNA | 1646-1666 | yes | LRA, WB | 22451425 |
| Tia1 | RBP | 1426-1442, 1702-1731 | yes, no | LRA, iCLIP | 28904350 |
| Hzf | RBP | 1345-1395, 1529-1574 | yes, yes | LRA, EMSA, WB, IP, RT-qPCR, Polysome gradient | 21402775 |

Abbreviations: LRA: luciferase reporter assay; WB: western blot; IP: co-immunoprecipitation assay; RT-qPCR: quantitative reverse transcription PCR; NB: northern blot; IHC: immunohistochemistry; ISH: In situ hybridization; EMSA: electromobility shift assay.

Here, we generated human cell lines and mice using CRISPR/Cas9 to delete the *TP53* and *Trp53* 3′UTRs at orthologous human and mouse gene loci while keeping mRNA processing intact. In HCT116 cells and in mouse tissues, we did not observe 3′UTR-dependent differences in p53 mRNA or protein levels under normal conditions or after DNA damage. When using the *TP53* 3′UTR in isolation, we confirmed the previously observed repressive effects in reporter assays. However, adding the p53 coding region to the reporters had a substantially stronger repressive effect on expression than the 3′UTR. Moreover, the presence of the p53 coding region prevented repression by the 3′UTR.

## Results

### Removal of the endogenous 3′UTR does not alter p53 mRNA or protein expression

3′UTRs perform two general functions: They contain regulatory elements that enable mRNA 3′ end processing and they harbor elements that allow post-transcriptional gene regulation (Matoulkova et al., 2012). 3′ end processing is essential for the generation of mature mRNAs and is facilitated by the poly(A) signal together with surrounding sequence elements that bind the polyadenylation machinery (Martin et al., 2012). Based on the binding motifs of polyadenylation factors, we consider 100-150 nucleotides upstream of the cleavage site as essential (Martin et al., 2012). Because the human *TP53* 3′UTR has a total length of about 1,200 nucleotides, the additional sequence could enable regulatory functions mediated by miRNAs and RNA-binding proteins. Indeed, the vast majority of previously characterized binding sites for miRNAs and RNA-binding proteins are located in the upstream, non-essential part of the *TP53* 3′UTR (Figure 1a, Table 1).

**Figure 1.**
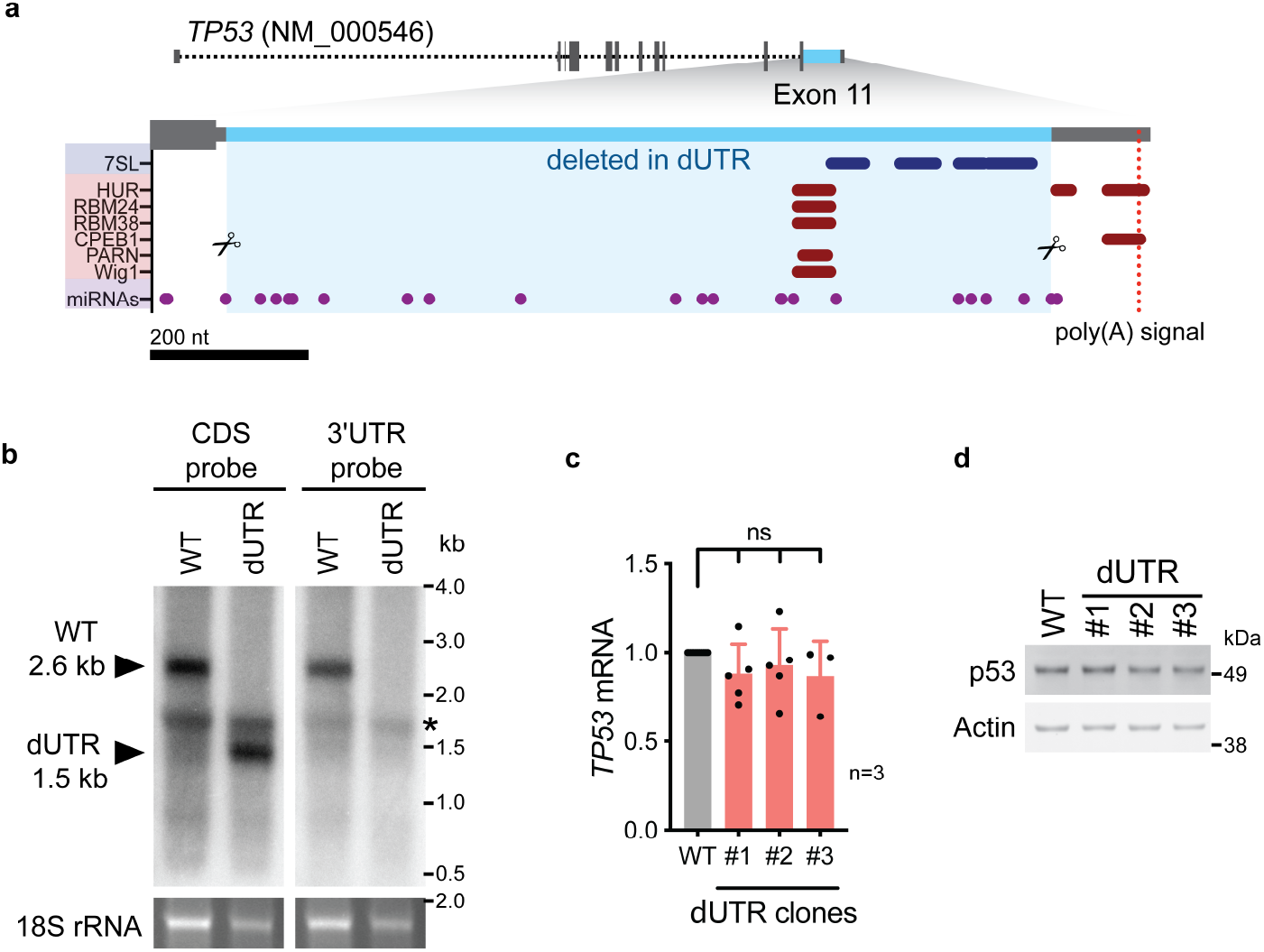
3’UTR-independent p53 expression regulation in steady state in human cells. **a,** Schematic of the human *TP53* gene. The sequence deleted in dUTR cells is shown in blue. Tracks of binding sites for miRNAs, RBPs and lncRNA are depicted below (see also Table 1). **b,** Northern blot analysis of *TP53* mRNA from WT and dUTR HEK293 cells. A probe that hybridizes to the *TP53* coding region (CDS) reveals expression of a shortened *TP53* mRNA in dUTR cells. The size difference is consistent with the length of the CRISPR/Cas9-induced deletion. A probe designed to bind the *TP53* 3’UTR does not produce a signal in the mRNA of dUTR cells, confirming removal of this sequence element. The band of 18S rRNA is used as a loading control. * indicates an unspecific band from ribosomal rRNA. **c,***TP53* mRNA expression levels in WT HCT116 cells and three different dUTR cell clones are shown from n=3 independent experiments (mean+ s.d.) after normalization to *GAPDH.* **d,** As in c, but shown is p53 protein expression. Actin serves as loading control. See also figure supplement 1 for sequnce alignments and data generated with *TP53* dUTR HEK293 cells.

To investigate the role of the endogenous human *TP53* 3′UTR in post-transcriptional p53 regulation, we used a pair of CRISPR/Cas9 guide RNAs to delete the non-essential part of the 3′UTR in HEK293 cells and in the human colon carcinoma cell line HCT116, an established model for investigating p53-dependent functions (Figure 1a, blue and Figure 1-supplement 1a). The homozygous 3′UTR deletion, called ΔUTR (dUTR), removed 1,048 nucleotides, corresponding to 88% of the 3′UTR in wild-type (WT) cells. The deletion affected almost all previously reported binding sites for regulatory miRNAs, lncRNAs, and RNA-binding proteins (Figure 1a, Table 1). We confirmed intact 3′ end processing of the mRNA by northern blot analysis and observed expression of the expected shorter *TP53* mRNA in dUTR cells (Figure 1b). We analyzed several HCT116 cell clones carrying a homozygous deletion of the *TP53* 3′UTR for p53 mRNA and protein expression and did not observe any differences in steady state cultivation conditions (Figures 1c and 1d). The same was true for HEK293 cells carrying the homozygous dUTR deletion (Figure 1-supplements 1b and 1c).

### The endogenous 3′UTR is not involved in regulating p53 levels after stress

While p53 mRNA expression does not change upon DNA damage, upregulation of p53 protein expression is achieved through higher translation rates and lower protein turnover (Kumari et al., 2014). Previous studies had suggested a role of the 3′UTR in the upregulation of p53 translation after exposure to genotoxic stress (Fu and Benchimol, 1997; Mazan-Mamczarz et al., 2003; Chen and Kastan, 2010). To assess stress-induced p53 expression regulation in dUTR cells, we treated cells with the topoisomerase inhibitor etoposide. We found that concentration-dependent upregulation of p53 protein expression was equal in WT and dUTR cells (Figure 2a). In addition, p53 levels analyzed over two days revealed similar p53 expression kinetics (Figure 2b). Finally, we tested additional stress stimuli including Nutlin-3 (an inhibitor of MDM2), 5-fluorouracil (a thymidylate synthase inhibitor) or UV irradiation. All of these treatments resulted in robust upregulation of p53 protein, but with no detectable differences in p53 expression between WT and dUTR cells (Figure 2c). We therefore concluded that the endogenous p53 3′UTR is not required for p53 expression regulation either in steady state or after DNA damage.

**Figure 2.**
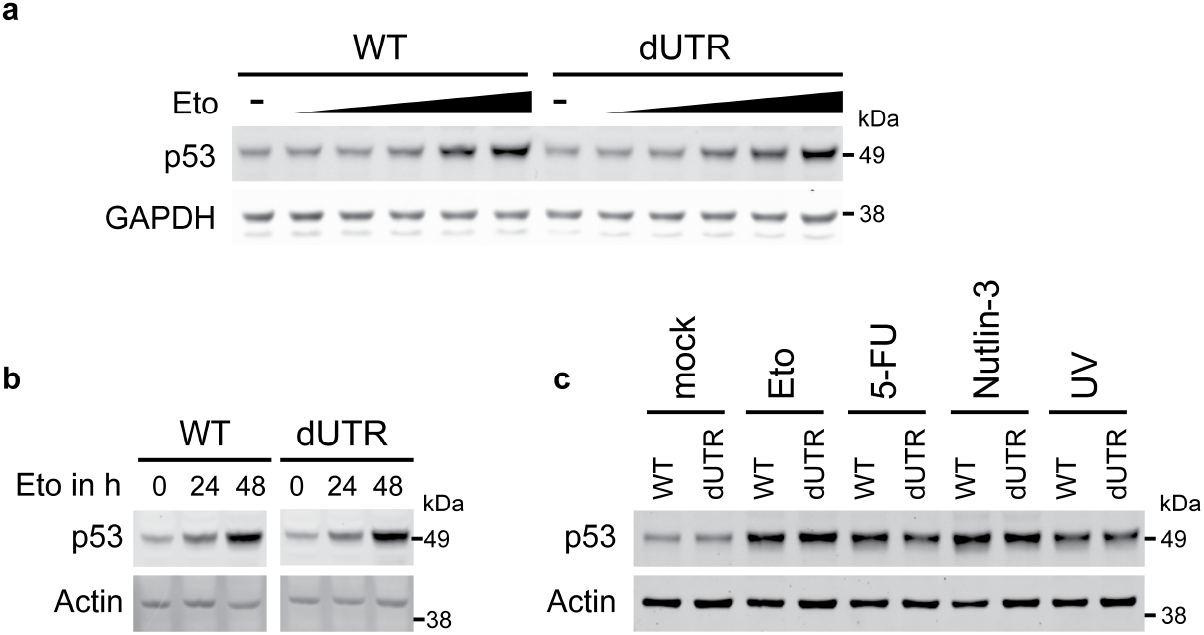
The 3’UTR is not required to induce upregulation of p53 protein after genotoxic stress. **a**, lmmunoblot showing p53 protein levels after four hours of Etoposide (Eto) treatment (0-32 μM) in WT and dUTR HCT116 cells. GAPDH serves as loading control. **b,** WT and dUTR HCT116 ells were treated with 0.5 μM Etoposide for 0, 24 and 48 hours. Actin serves as loading control. **c**, As in b, but cells were treated with 20 μM Etoposide, 40 μM 5-Fluorouridine (5-FU), 20 μM Nutlin-3 or 50 J/m^2^ UV. Actin serves as loading control.

### 3′UTR-mediated effects on reporter gene expression are context-dependent

We tried to reconcile our own findings using a genetic model with the existing studies suggesting a repressive function of the 3′UTR. Notably, earlier studies that investigated 3′UTR-dependent p53 regulation used reporter genes as proxy for endogenous p53 regulation (Table 1). We therefore cloned the human *TP53* 3′UTR (1,207 nucleotides) or the dUTR fragment (157 nucleotides) downstream of GFP and expressed these constructs in p53−/−HCT116 cells (Figure 3a, Figure 3-figure supplement 1a). In the context of the reporter, the *TP53* 3′UTR significantly reduced expression of both GFP mRNA and protein (Figures 3b and 3c). This result was recapitulated when luciferase was used instead of GFP reporters, thus confirming previous findings (Figure 3-figure supplement 1b). We wondered whether the endogenous sequence context could explain these discrepancies and added the p53 coding region (CDS) to our reporter constructs. As expected, we found that the CDS-GFP fusion protein was expressed at much lower levels than GFP alone, which could be due to high p53 turnover caused by MDM2. Surprisingly though, the p53 CDS also drastically suppressed expression of the reporter mRNA indicating a strong contribution of the CDS to p53 mRNA stability regulation (Figure 3c). Importantly, addition of the *TP53* 3′UTR in the context of the CDS did not further repress mRNA or protein expression of the GFP reporter, thus abrogating the difference between the samples containing the dUTR or full-length 3′UTR (Figure 3c). These results reveal that the *TP53* 3′UTR and CDS functionally interact in the regulation of p53 expression and that individual effects cannot be assumed to be additive.

**Figure 3.**
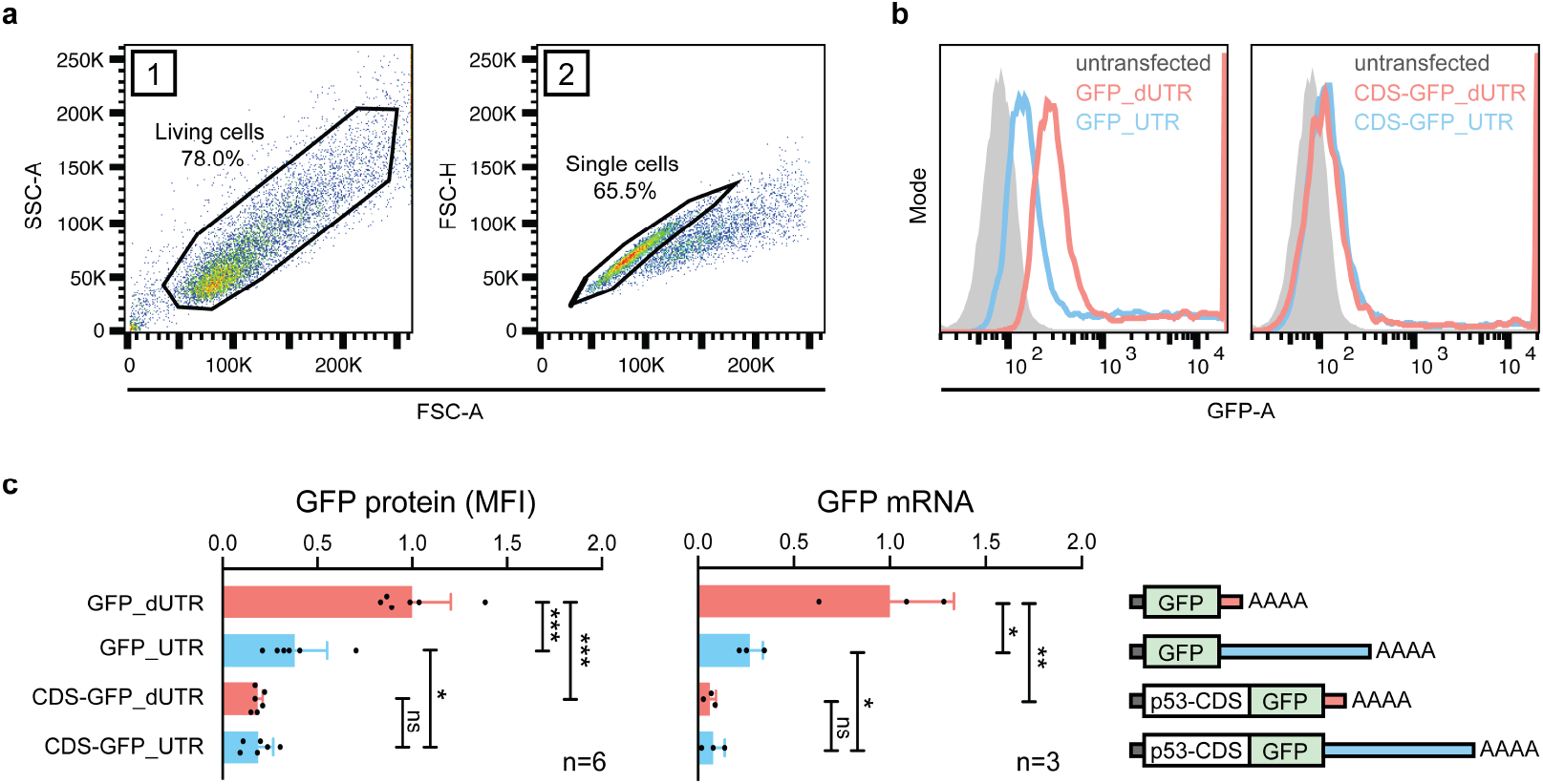
3’UTR and coding region of p53 have non-additive effects on the expression of a reporter gene. **a,** Gating strategy in FACS experiment for measurement of GFP protein expression in p53 −/− HCT116 cells. **b,** Histogram plots from one representative FACS experiment. The grey area represents the untransfected, GFP-negative control population. **c**, GFP protein levels were quantified by FACS and *GFP* mRNA levels were measured by RT-qPCR using *GAPDH* as housekeeping gene in p53 −/− HCT116 cells. Shown is mean + s.d. of n=3 independent experiments. CDS, coding sequence. Statistical analysis using unpaired Student’s t-test with * p<0.05, ** p<0.01, *** p<0.0001; ns not significant. See also figure supplement 1 for additional information.

### A Trp53 dUTR mouse model reveals 3′UTR-independent p53 expression *in vivo*

We reasoned that 3′UTR-dependent p53 expression regulation might still play a role in certain developmental stages, tissues or cell types. In order to explore this possibility, we created an analogous mouse model to investigate the role of the p53 3′UTR in an organism. We used zygotic injection of a pair of CRISPR/Cas9 guide RNAs to create mice in which we deleted the non-essential part of the mouse *Trp53* 3′UTR (Figure 4a). After backcrossing, we analyzed *Trp53* dUTR mice harboring a homozygous 3′UTR deletion (Figure 4-supplements 1a-c). These mice were viable, fertile, and did not show any developmental defects (Figure 4-supplements 1d and 1e). We measured *Trp53* mRNA expression in ten different tissues and did not detect significant differences between samples derived from WT and dUTR mice (Figure 4b). To examine the role of the 3′UTR in the regulation of stimulus-dependent p53 expression, we performed total body irradiation of WT and dUTR mice. At four hours post-irradiation, p53 protein expression was upregulated to a similar extent in spleen, liver, and colon samples from WT and dUTR mice (Figure 4c). We also analyzed expression of *Cdkn1a,* a highly dosage-sensitive p53 target gene that encodes the cell cycle regulator p21 (Fischer, 2019). Four hours after irradiation, *Cdkn1a* mRNA level were equally induced in WT and dUTR mice, suggesting that p53 target gene activation is 3′UTR-independent in mouse tissues (Figure 4d). These results demonstrate that the non-essential part of the *Trp53* 3′UTR is not required for steady state or stimulus-dependent regulation of p53 mRNA or protein level in mice.

**Figure 4.**
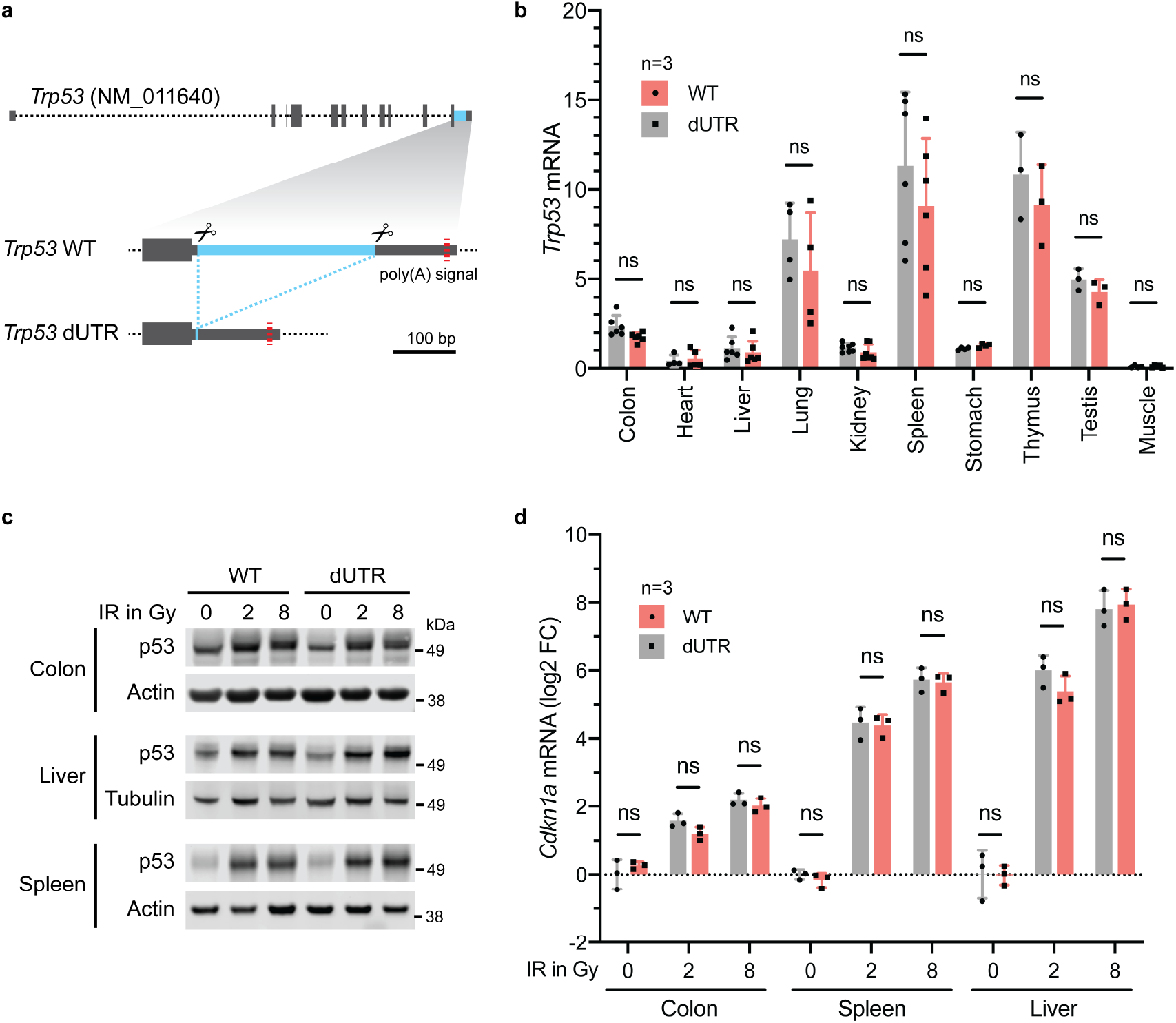
Knockout of the *Trp53* 3’UTR does not lead to aberrant p53 expression in a mouse model. **a,** Schematic of the mouse *Trp53* gene. The sequence deleted in dUTR cells is shown in blue. **b,***Trp53* mRNA in tissues from WT and dUTR mice was normalized to *Gapdh.* Shown is mean + s.d. from n= 3 independent experiments. See also figure supplement 1 for more information. **c,** Representative immunoblots of p53 protein from tissues obtained four hours after total body irradiation. Gy, Gray. **d,***Cdkn1a* mRNA expression of samples from (c) was normalized to *Gapdh.* Shown is mean + s.d. from three mice.

## Discussion

3′UTRs play important roles in the regulation of mRNA and protein abundance as well as in specifying protein functions (Mayr, 2019). A number of studies have previously proposed that the p53 3′UTR may be required to maintain low expression levels of p53 in non-stressed conditions (Haronikova et al., 2019). Especially miRNAs targeting p53 were previously established as putative gatekeepers to prevent p53 hyperactivation. In addition, some of these miRNAs are also elevated in cancer, e.g. miR-504, miR-30d, and miR-125 (Hu et al., 2010; Li et al., 2012; Banzhaf-Strathmann and Edbauer, 2014). This has sparked an interest in exploiting these mechanisms for therapeutic applications to modulate p53 expression level using novel miRNA-based approaches (Kasinski and Slack, 2011; Hermeking, 2012).

The lack of experimental data for 3′UTR-mediated expression regulation in native gene contexts has been a longstanding problem in the field of post-transcriptional gene regulation. Until recently, research on 3′UTR functions has mostly been conducted using overexpression systems and reporter gene assays. In contrast, gene knockouts that disrupt proteins have long been considered the gold standard for analyzing gene functions. The advent of CRISPR/Cas9 gene editing tools has made the creation of 3′UTR knockouts using genomic deletion feasible in both cell lines and organisms.

Using these tools, we observed that the endogenous p53 3′UTR does not have a significant impact on p53 abundance regulation. While we could reproduce earlier reporter studies with regards to a repressive function of the 3′UTR in isolation, we found that the 3′UTR-mediated repressive effect was abrogated in the context of the p53 coding region. This phenomenon may be explained by differences in RNA folding which could create constraints on motif accessibility. Our results indicate that the different parts of mRNAs do not act autonomously, but are part of a regulatory unit and functionally cooperate with each other (Cottrell et al., 2017; Theil et al., 2019). Notably, a recent study that deleted 3′UTR sequences in several cytokine genes found similar discrepancies between reporter-based assays and gene expression from native contexts (Zhao et al., 2017). Although our data indicate that p53 abundance regulation is 3′UTR-independent, the 3′UTR may still have important functions possibly through control of protein localization or protein activity as has been shown for other proteins (Berkovits and Mayr, 2015; Moretti et al., 2015; Terenzio et al., 2018; Lee and Mayr, 2019; Fernandes and Buchan, 2020; Bae et al., 2020; Kwon, 2020; Mayr, 2019).

Our observations further support the recently established role of the coding region as a major regulator of mRNA stability and translation (Mauger et al., 2019; Wu et al., 2019; Narula et al., 2019). Genome-wide comparisons of human coding regions showed that codon optimality and RNA structure in coding regions have the potential to modulate mRNA stability and translation efficiency to a similar extent as 3′UTRs.

RNA-binding proteins and miRNAs often target several members of a pathway (Ben-Hamo and Efroni, 2015; Zanzoni et al., 2019). Therefore, the results of overexpression or knockdown experiments of putative 3′UTR regulators may be confounded by other targets that might cause indirect effects. This issue might have contributed to the hypothesis of direct 3′UTR-dependent p53 regulation. For example, the tumor suppressor RBM38 (RNPC1) was proposed to bind to the human *TP53* 3′UTR resulting in lower p53 expression in the presence of RBM38 (Zhang et al., 2011). However, apart from p53, RBM38 targets several other genes in the p53 pathway, including *MDM2*, *PPMID*, and *CDKN1A* (Xu et al., 2013; Zhang et al., 2015; Shu et al., 2006). Expression changes of these genes can indirectly cause p53 expression regulation or result in phenotypes that mimic p53 overexpression. Indeed, while RBM38 knockout mice show phenotypes consistent with p53 hyperactivation (Zhang et al., 2014), Trp53 dUTR mice are apparently normal. This suggests that the repressive effects on p53 that were previously attributed to be mediated by 3′UTR-dependent abundance regulation may be indirect events.

Our data imply that in order to develop useful approaches for therapeutic intervention targeting post-transcriptional expression regulation, we need to develop a better understanding of these multi-layered regulatory networks. Our study shows that genetic manipulation of endogenous 3′UTRs may be a vital tool to disentangle direct from indirect post-transcriptional effects. It should become an essential step during the testing of miRNA-based therapies that are currently being explored as anti-cancer therapeutics in the context of p53 to avoid mixed or negative results in large clinical trials (Kasinski and Slack, 2011; Hermeking, 2012; Bonneau et al., 2019).

## Methods

### Generation of the *Trp53* dUTR mouse strain using CRISPR/Cas9

Female C57Bl/6 mice between 3-4 weeks of age were superovulated by intraperitoneal injection of Gestyl followed by human chorionic gonadotropin according to standard procedures (Behringer, 2014). After superovulation, the females were setup with male studs for mating. After mating, fertilized eggs were recovered at the one-cell stage from oviducts of superovulated female mice. 1-2 pl of CRISPR/Cas9 RNP complexes were injected into the pronuclei of fertilized eggs (see details below). Surviving eggs were surgically reimplanted into the oviducts of pseudo-pregnant females previously primed for pregnancy by mating with vasectomized males. The resulting pubs were screened using PCR for the deletion amplicon at two weeks of age (primers are listed in Supplementary Table 1). Suitable candidates were further validated by sequencing.

#### Preparation of CRISPR-Cas9 RNP injection mixture

Two target-specific crRNAs and a tracrRNA were purchased from IDT (Supplementary Table 1). In two separate tubes, 2.5 μg of each crRNA was mixed with 5 μg tracrRNA, heated to 95 °C for 5 min and then slowly cooled down to room temperature for annealing. The annealed duplexes were combined and mixed with 1 μg recombinant Cas9 enzyme (PNABIO) and 625 ng *in vitro* transcribed Cas9 mRNA and the total volume was adjusted to 50 μl with sterile water.

#### Screening for homozygous and heterozygous dUTR mice

Two heterozygous founder males with an identical 295 nucleotide deletion (Figure 4-figure supplement 1c) were used to establish a mouse colony. Two or more rounds of backcrossing into wildtype C57Bl/6 mice were performed prior to analysis of *Trp53* dUTR mouse phenotypes. Mouse genotypes from tail biopsies were determined using RT-PCR with specific probes designed for each Trp53 allele (Transnetyx, Cordova, TN).

#### Irradiation of mice

Where indicated, adult mice underwent total body irradiation with 2 or 8 Gy using a Cs-137 source in a Gammacell 40 Exactor (MDS Nordion) at 77 cGy/min. Four hours later irradiated mice were euthanized to collect samples. All procedures were approved by the Institutional Animal Care and Use Committee at MSKCC under protocol 18-07-010.

### Extraction of total RNA from mouse tissues and human cells for RT-qPCR analysis

For RNA extraction from mouse tissue, freshly collected tissue samples were flash-frozen and transferred to RNAlater-ICE Frozen Tissue Transition Solution (Invitrogen). After soaking overnight at −20 °C, the tissue samples were homogenized in vials containing 1.4 mm ceramic beads (Fisherbrand) and 400 μl RLT buffer (Qiagen) using a bead mill (Bead Ruptor 24, Biotage). 200 μl of the tissue homogenate was mixed with 1 ml of TRI Reagent (Invitrogen). For extraction of RNA from cultured cells, the cell pellet was directly resuspended in TRI Reagent. Total RNA extraction was performed according to the manufacturer’s protocol. The resulting RNA was treated with 2U DNaseI enzyme (NEB) for 30 min at 37 °C, followed by acidic phenol extraction and isopropanol precipitation. To generate cDNA, about 200 ng of RNA was used in a reverse transcription reaction with SuperScript IV VILO Master Mix (Invitrogen). To measure the relative expression levels of mRNAs by RT-qPCR, FastStart Universal SYBR Green Master (ROX) from Roche was used together with gene-specific primers listed in Supplementary Table 1. GAPDH/Gapdh was used as reference gene.

### Generation of the *TP53* 3′UTR deletion in HCT116 and HEK293 cells

To generate CRISPR/Cas9 constructs, we annealed target-specific gRNA sequences and inserted them into a BbsI-digested pX330-U6-Chimeric_BB-CBh-hSpCAs9 vector (Addgene plasmid #42230) (Cong et al., 2013; Ran et al., 2013). 1 μg of each pX330-gRNA plasmid plus 0.1 μg of pmaxGFP plasmid (Lonza) were transiently transfected into exponentially growing cells using Lipofectamine 2000 (Invitrogen). Three days after transfection, single GFP-positive cells were sorted into 96-well plates and cultured until colonies formed. The genomic DNA from individual cell clones was extracted using QuickExtract DNA Extraction Solution (Lucigen) and screened by PCR for the deletion amplicon using the DNA primers listed in Supplementary Table 1. In the case of HCT116 cells, we repeated the above-described process using two different heterozygous clones with a new downstream gRNA to obtain homozygous *TP53* dUTR cells. Finally, to validate positive cell clones, all *TP53* alleles of candidate clones were sequenced (Figure 1-supplement 1a).

### Generation of p53 KO HCT116 and HEK293 cells

We generated our own p53-deficient HEK293 and HCT116 cell lines by targeting exon 6 of the p53 coding region with a gRNA causing frame shift mutations. Specifically, pX330 plasmid harboring a p53-specific gRNA (Supplementary Table 1) was transfected into HEK293 and HCT116 cells using Lipofectamine 2000 (Invitrogen). Two days later, the cells were split and seeded sparsely on a 10 cm dish in the presence of 10 μM Nutlin-3 (Seleckchem) which was used to select against growth of p53-competent cells. After ten days, single colonies were picked, and individual clones were validated by WB for loss of p53 expression.

### Western blot analysis

RIPA buffer (10 mM Tris-HCL pH 7.5, 150 mM NaCl, 0.5 mM EDTA, 0.1% SDS, 1% Triton X-100, 1% deoxycholate, Halt Protease Inhibitor Cocktail (Thermo Scientific)) was used to extract total protein from cultured cells or mouse tissues. Cell pellets were washed with PBS and directly resuspended in lysis buffer and incubated on ice for 30 min. Mouse tissue samples were homogenized in RIPA buffer using a bead mill in vials filled with 1.4 mm ceramic beads. Tissue lysates were sonicated to shear genomic DNA prior to removing insoluble components by centrifugation (10 min, 15,000 g). The proteins in the supernatant were precipitated by adding 0.11 volumes of ice-cold 100 % Trichloroacetic acid (TCA) and incubated at −20 °C for one hour. The samples were centrifuged (10 min, 15,000 g) and the pellet was washed twice in ice-cold acetone before resuspending in reducing 2x Laemmli buffer (Alfa Aesar). Proteins were separated by size on a 4-12% Bis-Tris SDS-PAGE gels (Invitrogen) and blotted on a 0.2 μm nitrocellulose membrane (BIO-RAD). The membrane was then incubated with primary antibody in Odyssey Blocking buffer (LI-COR) overnight at 4 °C. The following primary antibodies were used in this study: anti-human p53 (Santa Cruz, sc-47698, mouse, 1:250), anti-mouse p53 (Cell Signaling, #2524, mouse, 1:500), anti-Actin (Sigma, A2008, rabbit, 1:1000), anti-Tubulin (Sigma, T9026, mouse 1:1000) and anti-GAPDH (Sigma, G8705, mouse, 1:1000). After washing, the membrane was incubated with fluorescently-labeled secondary antibodies (IRDye 800CW Goat anti-Mouse, 926-32210; IRDye 680 Goat anti-Rabbit, 926-68071 LI-COR) and signals were recorded using the Odyssey Infrared Imaging system (LI-COR).

### Northern Blot

Total RNA from cells was extracted as described above. Afterwards, polyA+ mRNA was enriched from total RNA using the Oligotex suspension (Qiagen) according to the manufacturer’s instructions. 1.2 μg of polyA+ mRNA was glyoxylated and run on an agarose gel as described previously (Mayr and Bartel, 2009). The RNA was transferred overnight using the Nytran SuPerCharge TurboBlotter system (Whatman) and UV-crosslinked.

DNA probes complementary to the *TP53* coding region or the 3′UTR were labeled with dCTP [α-^32^P] using the Amersham Megaprime DNA labeling system (GE Healthcare). Primers used for probe synthesis from human cDNA are listed in Supplementary Table 1. Labeled probes were denatured by heat for 5 min at 90 °C and then incubated with the blot in ULTRAhyp Ultrasensitive Hybridization Buffer (Invitrogen) overnight at 42 °C. The blot was washed three times and exposed on a phosphorimaging screen. The radioactive signal was acquired using the Fujifilm FLA700 phosphorimager.

### Human cell culture and drug treatment

Human cell cultures were maintained in a 5% CO2/ 37 °C humidified environment. HEK293 cells were cultured in DMEM (high glucose) and HCT116 cells were cultured in McCoy’s 5A medium which were supplemented with 10% FBS and 1% Penicillin/Streptomycin. Where indicated, HCT116 cells were treated with etoposide (0.125-32 μM, Sigma), 5-fluorouracil (40 μM, Sigma), Nutlin-3 (20 μM, Seleckchem), or UV (50 J/m^2^) prior to downstream analysis.

### Reporter assays

We PCR-amplified the *TP53* 3′UTR sequence (nucleotides 1,380 to 2,586 of the reference mRNA NM_000546, May 2018) from WT HCT116 cDNA. This sequence was cloned downstream of the stop codon in pcDNA3.1-puro-eGFP using EcoRI/NotI restriction enzymes. For the dUTR construct, cDNA from TP53 dUTR HCT116 cells was used to amplify the remaining 3′UTR sequence after CRISPR-mediated deletion, representing a fusion of the first 12 and the last 157 nucleotides of the *TP53* 3′UTR. The p53 coding region, encoding the α protein isoform (1,182 nucleotides), was cloned upstream and in frame of the GFP-cassette using HindIII/BamHI restriction sites. For luciferase reporter studies, the full length 3′UTR and dUTR sequences described above were cloned into a SmaI-digested psiCHECK2 (Promega) vector via blunt-end cloning.

#### GFP reporter

GFP protein levels of cells transfected with equimolar amounts of GFP-containing reporter constructs was analyzed by flow cytometry after 24 hours. A BD LSRFortessa Flow Cytometer was used to record the mean fluorescence intensity (MFI) of 20,000 life cells. Raw data were analyzed using the FlowJo software package and values were normalized to GFP-only constructs. mRNA abundance of the GFP reporter was measured using RT-qPCR using the primers listed in Supplementary Table 1. The GFP reporter mRNA was normalized to *GAPDH* mRNA.

#### Luciferase reporter assay

Luciferase activity was measured 24 hours after transfection of equimolar amounts of psiCHECK2 plasmids (Promega) containing either the *TP53* 3′UTR or dUTR sequence downstream of the Renilla luciferase translational stop codon. Cells were lysed in passive lysis buffer and Renilla and firefly luciferase activity was measured in duplicates using the Dual-Glo Luciferase Assay System (Promega) according to the manufacturer’s instructions in a GloMax 96 Microplate Luminometer (Promega). Relative light units of Renilla luciferase were normalized to firefly luciferase activity.

### Statistics and reproducibility

Statistical analysis of the mRNA and protein expression data was performed using a Student’s t-test or ANOVA followed by a Tukey’s multiple comparison test. We use ns (*p* > 0.05), * 0.01 < *p* < 0.05, ** 0.001 < *p* < 0.01, and *** *p* < 0.001 to indicate the levels of *p-*values in figures. No data were excluded. The results for immunoblotting are representative of at least three biologically independent experiments. All statistical analyses and visualizations were performed by using GraphPad (Prism 8).

## Acknowledgements

We thank all members of the Mayr lab for helpful discussions and critical reading of the manuscript. We thank the Mouse Genetics Core Facility at MSKCC for assistance in the generation of Trp53 dUTR mice. This work was funded by a postdoctoral fellowship from the DFG to S.M. and by the NIH Director′s Pioneer Award (DP1-GM123454), the Pershing Square Sohn Cancer Research Alliance to C.M., and the NCI Cancer Center Support Grant (P30 CA008748). The funders had no role in study design, data collection and interpretation, or the decision to submit the work for publication.

## Author contributions

S.M. performed all experiments and analyses. S.M. and C.M. conceived the project, designed the experiments, and wrote the manuscript.

## Declaration of Interests

The authors declare no competing interests.

## Additional files

Supplementary Table 1. Primer sequences

Transparent Reporting Form

## Data availability

All data generated and analyzed are included in the manuscript and supporting files.

---

**Figure 1-figure supplement 1.**
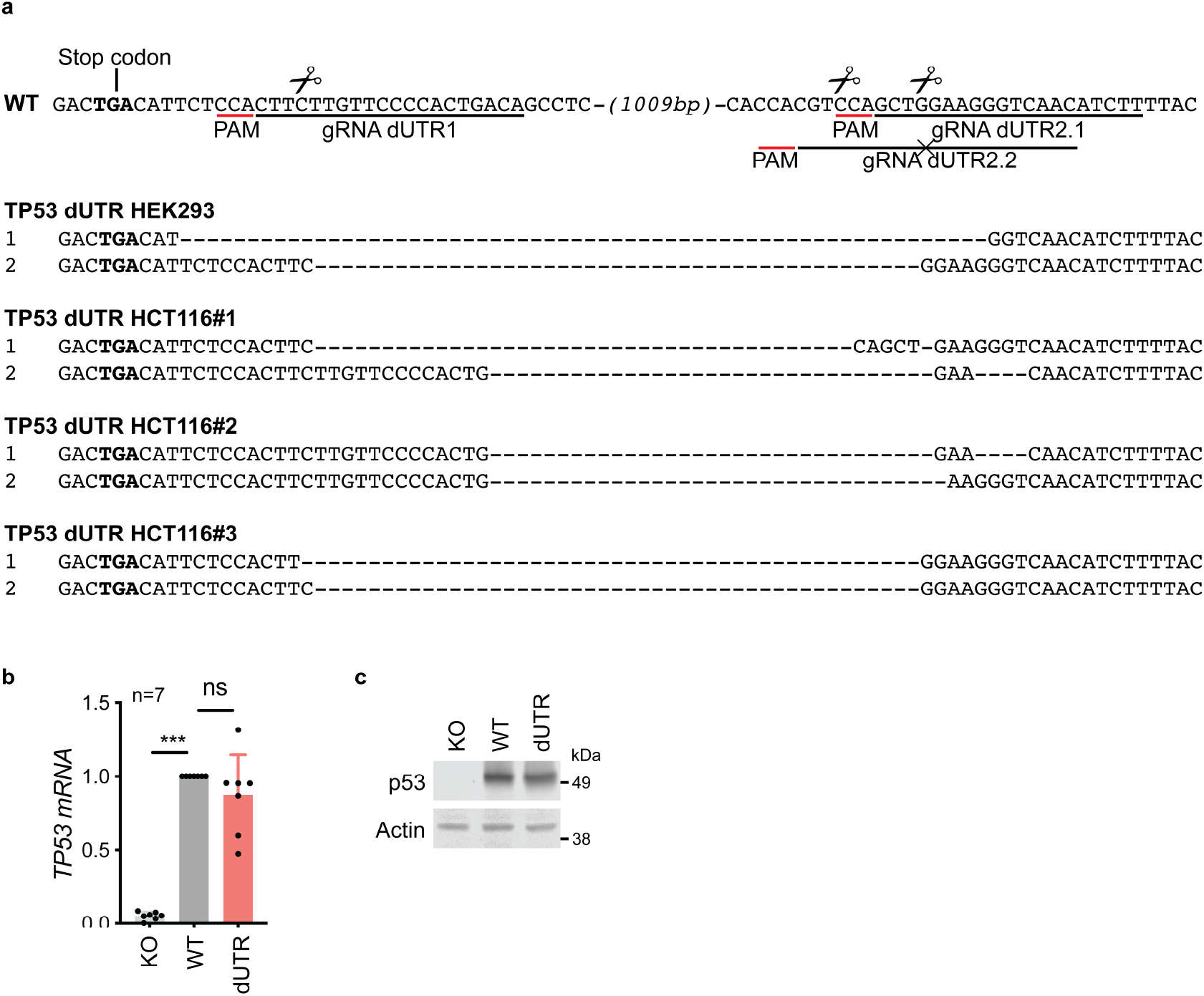
Generation and characterization of *TP53* dUTR human cell lines. **a,** Sequence alignment of TP53 alleles spanning the deletion sites in WT and dUTR HEK293 and HCT116 cell clones analyzed in this study. Binding sites of gRNAs used to generate the deletion are underlined in the WT reference sequence and predicted cutting sites are marked by a scissor symbol. gRNA dUTR2.2 harboring a specifc point mutation relative to the WT allele was used to create homozygous dUTR HCT116 cell lines during a second round of transfection. **b,** Analysis *TP53* mRNA levels in WT, protein KO, and dUTR HEK293 cells was measured by RT-qPCR and normalized with *GAPDH* mRNA levels. Shown are mean + s.d. of n= 7 indepen dent experiments. **c,** lmmunoblot showing p53 protein level in HEK293 cells, grown under steady state conditions.

**Figure 3-figure supplement 1.**
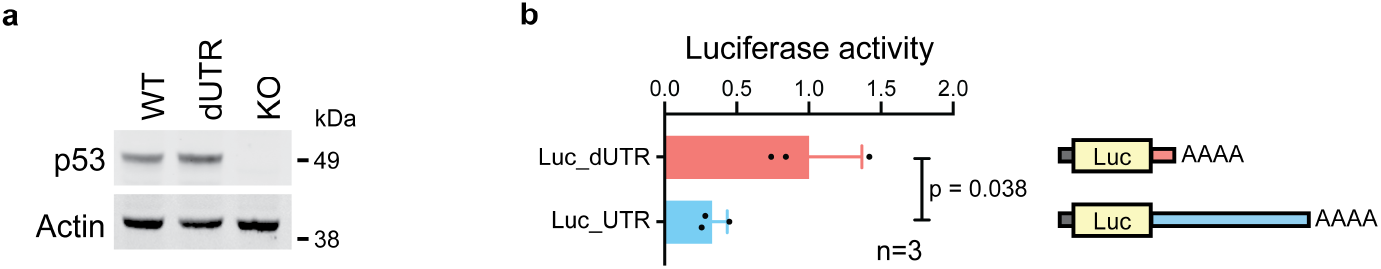
Validation of repressive effect of *TP53* 3’UTR in luciferase reporter assay. **a,** lmmunoblot showing p53 protein level in WT, dUTR and p53−/− HCT116 cells, grown under steady-state conditions. **b,** Renilla luciferase activity of constructs containing either the human dUTR or the human full-length *TP53* 3’UTR was performed in p53 −/− HCT116 cells. Shown is mean+ s.d. of n=3 independent experiments after normalization to firefly luciferase. Statistical analysis using t-test for independent samples.

**Figure 4-figure supplement 1.**
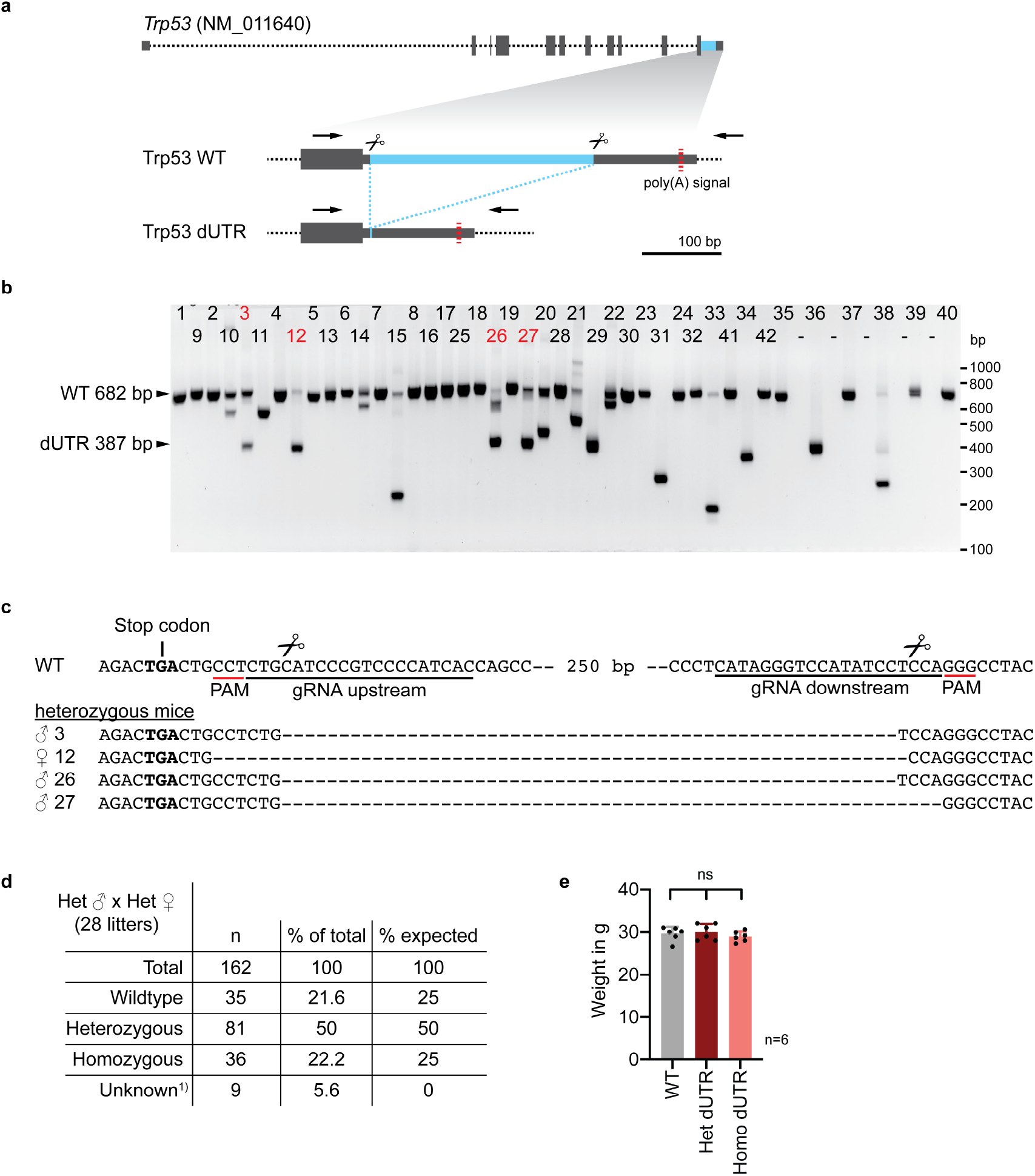
Generation and characterization of *Trp53* dUTR mice. **a,** Schematic of the mouse *Trp53* gene. The sequence deleted in dUTR cells is shown in blue and binding sites of primers used for PCR screening are marked with arrows. **b,** Screening PCR of mice that were born after zygotic injection of CRISPR/Cas9 RNPs targeting the *Trp53* 3’UTR. The predicted lengths of the PCR products from WT and dUTR alleles are indicated. Mice that were selected for validation by sequencing are labeled in red. **c,** Sequence alignments of *Trp53* dUTR alleles of select founder mice shown in b. Male mice #3 and #26 harboring identical DNA deletions were used to establish a mouse colony. Primer sequences used for screening can be found in Supplementary Table 1. **d,** Genotypes of pups from 28 *Trp53* dUTR heterozygous intercrosses. Unknown refers to mice that died before weening. **e,** Weights of mice at 10-11 weeks of age are shown for WT, *Trp53* dUTR heterozygous and homozygous males. Data are shown as mean + s.d. with n=6 in each group.

## Notes

### Competing Interest Statement

The authors have declared no competing interest.

### Summary of Updates

We have edited the text of the manuscript for more clarity. Furthermore, we have rearranged the figure panels to enable a more intuitive structure of arguments. However, we did not add or remove any data from the manuscript or alter the interpretation of the data.

